# Rapid oxygen titration following cardiopulmonary resuscitation mitigates cerebral overperfusion and striatal mitochondrial dysfunction in asphyxiated newborn lambs

**DOI:** 10.1101/2024.07.27.603805

**Authors:** Shiraz Badurdeen, Robert Galinsky, Calum T. Roberts, Kelly J. Crossley, Valerie A. Zahra, Alison Thiel, Yen Pham, Peter G. Davis, Stuart B. Hooper, Graeme R. Polglase, Emily J. Camm

## Abstract

Asphyxiated neonates must have oxygenation rapidly restored to limit ongoing hypoxic-ischemic injury. However, the effects of transient hyperoxia after return of spontaneous circulation (ROSC) are poorly understood. We randomly allocated acutely asphyxiated, near-term lambs to cardiopulmonary resuscitation in 100% oxygen (“standard oxygen”, n=8) or air (n=7) until 5 minutes after ROSC, or to resuscitation in 100% oxygen immediately weaned to air upon ROSC (“rapid-wean”, n=7). From 5 minutes post-ROSC, oxygen was titrated to target preductal oxygen saturation between 90-95%. Cerebral tissue oxygenation was transiently but markedly elevated following ROSC in the standard oxygen group compared to the air and rapid-wean groups. The air group had a delayed rise in cerebral tissue oxygenation from 5 minutes after ROSC coincident with up-titration of oxygen. These alterations in oxygen kinetics corresponded with similar overshoots in cerebral perfusion (pressure and flow), indicating a physiological mechanism. Transient cerebral tissue hyperoxia in the standard oxygen and air groups resulted in significant alterations in mitochondrial respiration and dynamics, relative to the rapid-wean group. Overall, rapid-wean of oxygen following ROSC preserved striatal mitochondrial respiratory function and reduced the expression of genes involved in free radical generation and apoptosis, suggesting a potential therapeutic strategy to limit cerebral reperfusion injury.

## Introduction

Cardiopulmonary resuscitation (CPR) of neonates with severe perinatal asphyxia currently focuses on rapid restoration of the circulation to limit ongoing hypoxic-ischemic brain injury.^1,2^ Given the challenges in conducting adequately powered trials of CPR in neonates, international recommendations rely on transitional animal models to guide practice.^1^ Current guidelines recommend increasing the fraction of inspired oxygen (FiO_2_) to 100% if chest compressions are required to restore myocardial contractility.^2,3^ Following return of spontaneous circulation (ROSC), the optimal strategy for oxygen provision is not known. In practice, oxygen is typically titrated to mimic oxygen saturation (SpO_2_) levels of neonates with normal perinatal transition.^2,4^ However, due to the rapid cerebral overperfusion that occurs immediately after ROSC, this strategy results in global cerebral oxygen delivery that is markedly in excess of oxygen consumption in the minutes following ROSC which can lead to oxidative stress and disruption of cerebral metabolism.^5–11^

The effect of transient excess oxygen delivery immediately after ROSC on region-specific cerebral tissue oxygenation and cellular function is unknown. Acute perinatal asphyxia predominantly affects the deep grey nuclei (striatum) and the perirolandic cortex.^12^ Following asphyxia and re-perfusion, alterations in mitochondrial bioenergetics and mitochondrial- derived reactive oxygen species (ROS) result in oxidative damage to mitochondrial proteins, membranes and DNA, impairing their ability to synthesize ATP, ultimately leading to cell death and secondary brain injury.^13,14^ Transient hyperoxia may therefore exacerbate ROS production, further impairing mitochondrial function and contributing to secondary energy failure that is characteristic of hypoxic-ischemic encephalopathy.^15^ Strategies to limit hypoxia while avoiding hyperoxia-mediated oxidative stress in the reperfusion phase have not been well studied.^8,16^

Among neonates with mild perinatal asphyxia requiring respiratory support alone, commencing mask ventilation in 100% oxygen versus air increases the risk of mortality.^17^ In neonates needing CPR therefore, the more severe initial insult may render pathways leading to secondary reperfusion injury particularly susceptible to even transient hyperoxic exposure. Here, we hypothesized that in acutely asphyxiated newborn lambs, a brief delay in weaning supplemental oxygen leads to cerebral tissue hyperoxia and mitochondrial dysfunction, which can be mitigated by rapidly weaning oxygen to air immediately after ROSC.

## Materials and methods

All procedures were approved by the Monash University Animal Ethics Committee (MMCA- 2020-04; 24/6/2020) and conducted in accordance with the National Health and Medical Research Council of Australia code for the care and use of animals for scientific purposes. This study is reported according to the ARRIVE guidelines.^18^ Additional methods are provided in the Supplementary Materials.

### Instrumentation

Border-Leicester sheep fetuses at 139±2 days’ gestation (mean ± SD; term ∼148 days) were intubated and instrumented after exteriorization of the head and chest via laparotomy. Cerebral oxygenation parameters were measured using fibre-optic oxygen probes placed in the right cerebral striatum (caudate nucleus) and right carotid artery. Carotid blood flow is known to strongly correlate with cerebral perfusion.^19^ Regional cortical tissue oxygen saturation (crSO_2_) was measured over the left frontal cortex using near-infrared spectroscopy (NIRS).

### Asphyxia and management after birth

Asphyxia was induced by cord clamping while withholding ventilation. At terminal asphyxia defined by asystole, mean arterial blood pressure of ∼0 mmHg and pH <7.0, ventilation was initiated in air with pressures of 30/5 cmH_2_O at 60 ventilations per minute (NeoPuff, Fisher & Paykel). One minute later, chest compressions were initiated at a 3:1 ratio with ventilation and FiO_2_ delivered according to one of 3 strategies that were randomly assigned prior to resuscitation using a web-based random sequence generator (www.random.org/lists):

1. Standard oxygen (100% oxygen from the commencement of chest compressions until 5 minutes after ROSC), n=8
2. Air (21% oxygen until 5 minutes after ROSC), n=7
3. Rapid-wean (100% oxygen from the commencement of chest compressions until onset of ROSC, upon which oxygen was immediately weaned to 21% until 5 minutes after ROSC), n=7.

Epinephrine (1:10,000, 0.2ml/kg) was administered 1 minute after initiating chest compressions. ROSC was defined as continuous unsupported cardiac output with diastolic blood pressure >20mmHg. Lambs then received ventilator-driven breaths at 30/5 cmH_2_O for 10 minutes followed by volume-guarantee mode (7 mL/kg). From 5 minutes post-ROSC, FiO_2_ was adjusted in all groups to target pre-ductal arterial oxygen saturation of 90-95%.

### Data Acquisition and Analysis

Continuous recordings of physiological parameters were made at 1 kHz. Carotid arterial blood gases were taken before induction of asphyxia, at terminal asphyxia, during CPR, at ROSC, and at regular intervals until 60 minutes after ROSC. All analyses were conducted blind to group allocation.

### Tissue Collection and Analysis

At 1 hour after ROSC, lambs were euthanized and tissue samples taken from the left striatum (matching the level where the oxygen probe was placed) and left cerebral cortex (at the level of the ansate sulcus, equivalent to the perirolandic cortex in humans). Mitochondrial respiratory rates and bioenergetics were measured using respirometry, western blotting and quantitative PCR (q-PCR), as previously described.^20,21^

### Statistical Analysis

From our previous work, 7-8 lambs per group were able to demonstrate differences in mitochondrial oxidative phosphorylation (OXPHOS).^21^ Baseline fetal characteristics, mitochondrial respirometry and western blot data were compared using a one-way analysis of variance, t-test or Mann-Whitney U test, as appropriate. Comparisons of physiological data between study groups were done using two-way repeated measures analysis of variance, or mixed effects models when there was missing data. Physiological data were analysed separately over the first 10 minutes, and then from 10-60 minutes. Tukey test was used to adjust p-values for multiple comparisons. Quantitative PCR data was expressed as relative change (2–ΔΔCT) were log-transformed to maintain normal distribution, and all statistical analysis and conducted on log-transformed 2–ΔΔCT data. Volcano plot data visualizations present comparisons of standard care versus rapid-wean or 21% oxygen as fold changes. P values were adjusted for multiple comparisons using the Benjamini-Hochberg method. Analysis was performed using PRISM version 9 for Mac OS X (GraphPad Software, La Jolla California USA).

## Results

### Baseline physiological comparisons

One lamb in the rapid-wean group died shortly after ROSC due to delayed recognition of a mal-positioned endotracheal tube and was excluded from all analyses. Baseline characteristics and blood gas parameters of the remaining 22 lambs were similar between groups except for fetal pO_2_, which was lowest in the standard oxygen group (Table 1). All lambs achieved ROSC following a single dose of epinephrine.

Table 1.

### Time to achieve ROSC was similar between lambs receiving CPR in 100% oxygen versus air

The mean time to achieve ROSC after commencing ventilation was similar between lambs receiving CPR in 100% oxygen (standard and rapid-wean groups, n=15, 185 s, 95% confidence interval [CI] 173–196 s) compared to the 21% oxygen group (n=7, 191 s, 95% CI 174–207 s). All lambs achieved ROSC following a single dose of epinephrine.

### Rapid-wean of FiO_2_ dampened the rise in cerebral tissue oxygenation following ROSC

During CPR, arterial oxygen saturation, striatal pO_2_ and crSO_2_ remained low despite the use of 100% oxygen in the standard and rapid-wean groups (Figures 1B-E). Upon ROSC, lambs in the air group had the lowest arterial oxygen saturation and crSO_2_ (Figures 1B, E). In the standard oxygen group, arterial oxygen saturation, carotid arterial pO_2_, striatal tissue pO_2_ and crSO_2_ were markedly elevated within 1-3 min post-ROSC (Figures 1B-E). In comparison, the fast rise in oxygenation was dampened in the rapid-wean group- for example, mean carotid arterial pO_2_ at 3 min post-ROSC was 56 mmHg vs 298 mmHg (*p* = 0.0007); striatal pO_2_ at 6 min post-ROSC was 29 mmHg vs 104 mmHg (*p* = 0.008). The air group had the slowest initial rise in oxygenation parameters.

**Figure 1.**
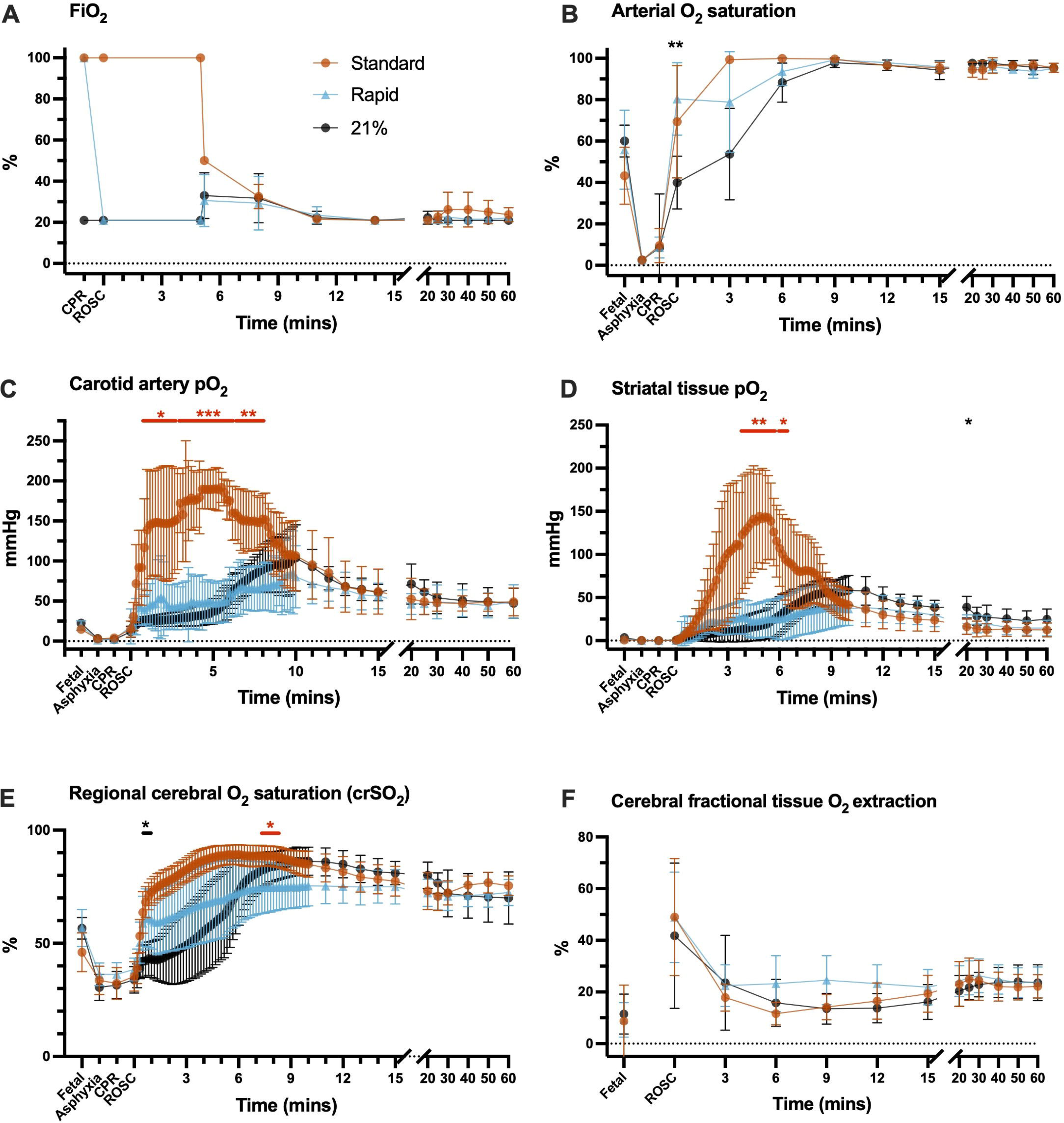
Fraction of inspired oxygen (FiO_2_, A) and oxygenation parameters (B-F) for lambs in the standard oxygen group (“Standard”, orange), rapid-wean group (“Rapid”, blue), and air group (“21%”, black). The standard oxygen group had a marked rise in carotid arterial oxygen saturation **(B)** and pO_2_ **(C)**, as well as striatal tissue pO_2_ **(D)** and regional cerebral oxygen saturation **(E)** from the onset of ROSC that was mitigated by rapidly weaning FiO_2_ to air. The air group had a slow initial rise in oxygenation parameters followed by a secondary rise once FiO_2_ was increased at 5 min after ROSC. Error bars represent 95% confidence intervals. Asterisks in red and black represent statistically significant differences between the rapid-wean versus standard oxygen group, and rapid-wean versus air groups, respectively. Comparisons between the standard oxygen and air group are not shown for clarity. * p <0.05, ** p <0.01, *** p <0.001; Tukey post hoc tests of mixed effects analysis.

From 5 minutes post-ROSC, all lambs had FiO_2_ titrated to target 90-95%. In the standard oxygen group, FiO_2_ was steadily weaned (Figure 1A). All lambs in the air group had FiO_2_ increased (range 25-50%) and 4/7 lambs in the rapid-wean group had FiO_2_ increased (range 30-50%). In the air group, this corresponded with a distinct secondary rise in striatal pO_2_ and crSO_2_ that was not apparent in the rapid-wean group, although differences did not meet statistical significance (Figures 1D, E). From approximately 12 minutes post-ROSC, oxygen parameters were similar between groups.

Regional cortical tissue oxygen saturation (Figure 1E) was significantly higher between 30 s – 5 min post-ROSC in the standard oxygen versus the air group (*p* < 0.05). From 5 min post-ROSC, the secondary rise in crSO_2_ that followed up-titration of FiO_2_ in the air group resulted in both crSO_2_ and cerebral fractional tissue oxygen extraction values that were similar to the standard oxygen group (Figures 1E, F). Correspondingly, peak crSO_2_ was ≥85% in 7/8 lambs and 6/7 lambs in the standard oxygen and air groups respectively. In contrast, 2/7 lambs in the rapid-wean group had peak crSO_2_ ≥85%.

### The post-ROSC overshoot in pulmonary and carotid blood flow matched cerebral oxygen kinetics and was least prominent in the rapid-wean group

Carotid arterial pO_2_ was significantly higher in the standard care group at 3 and 6 minutes after ROSC compared to the rapid wean and air groups (Figure 2A). Arterial pCO_2_ was similar between groups at all timepoints (Figure 2B). Pulmonary artery blood flow was higher in the standard care group for the first 5 minutes (by 21.3% vs. Rapid Wean and 21.0% vs. Air) but did not reach significance at individual time points (P_group x time_; *p* < 0.001; Figure 2C). Lambs in the air group had a delayed secondary rise in pulmonary artery blood flow following the increase in FiO_2_ from 5 min after ROSC, which was not apparent in the rapid-wean group.

**Figure 2.**
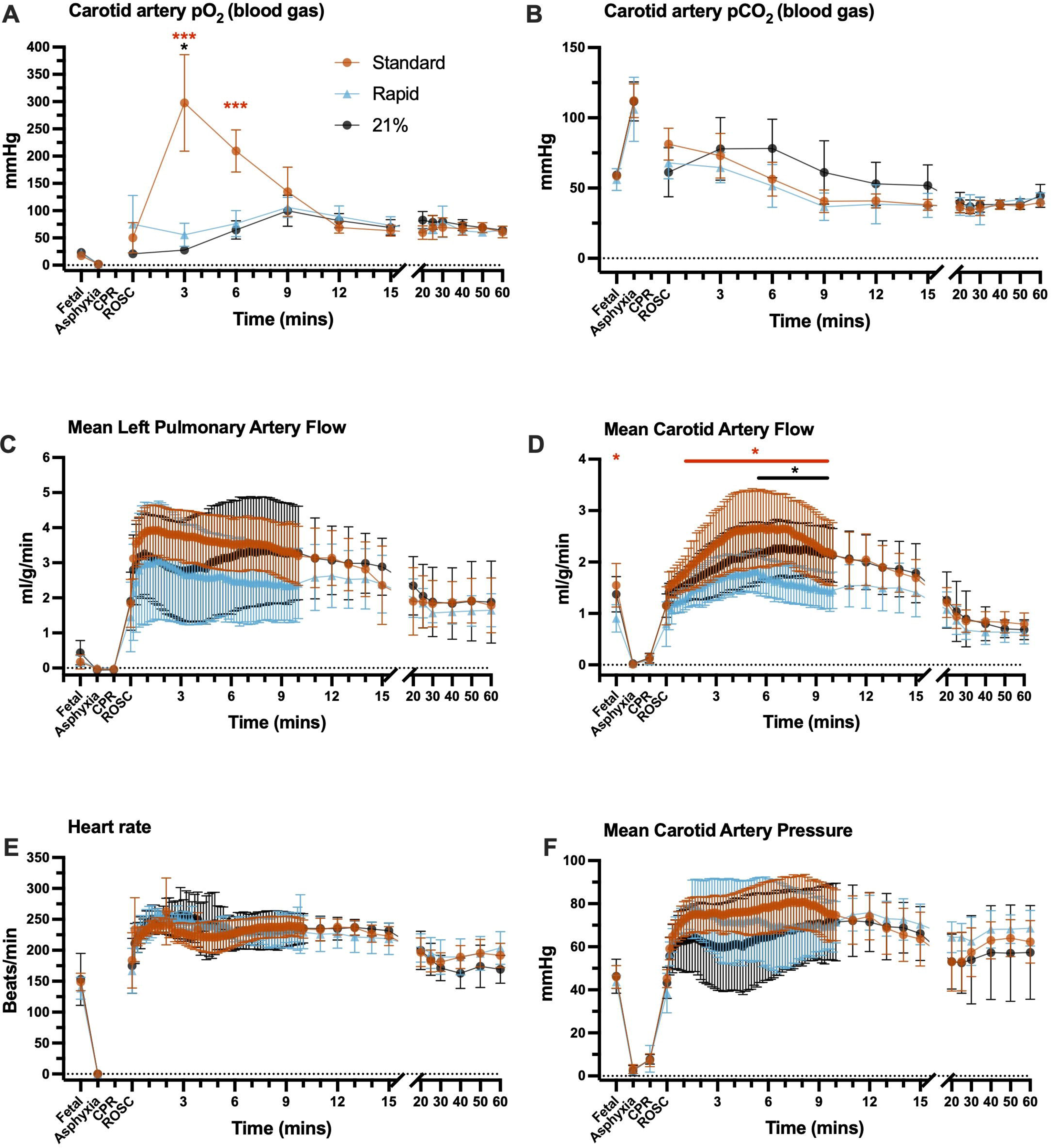
Blood gas pO_2_ (A), pCO_2_ (B) and cardiovascular parameters (C-F) for lambs in the standard oxygen group (“Standard”, orange), rapid-wean group (“Rapid”, blue), and air group (“21%”, black). The standard oxygen group tended to have the highest mean carotid artery blood pressure and the highest mean pulmonary artery blood flow in the first 5 minutes after ROSC, but differences between groups were not statistically significant. Lambs in the rapid-wean group had lower mean carotid blood flow both at the fetal timepoint and after ROSC when compared to the standard oxygen group. Error bars represent 95% confidence intervals. Asterisks in red and black represent statistically significant differences between the rapid-wean versus standard oxygen group, and rapid-wean versus air groups, respectively. Comparisons between the standard oxygen and air groups are not shown for clarity. * p <0.05, ** p <0.01, *** p <0.001; Tukey post hoc tests of mixed effects analysis.

Carotid arterial pressures were similar between groups (Figure 2F). However, the post-ROSC overshoot in carotid blood flow (Figure 2D) was lowest in the rapid-wean group, with a significantly lower carotid blood flow (P_group_ *p* = 0.025) than the standard care (from 20 s – 10 min; *p* < 0.05) and air (from 5:40 min – 10 min; *p* < 0.05) groups. Lambs in the standard care group had a rapid rise in both carotid arterial pressure and blood flow consistent with cerebral overperfusion, which was markedly reduced in the air and rapid-wean groups during the first 10 minutes. Taken together, the post-ROSC overshoot in pulmonary and carotid blood flows was least prominent in the rapid-wean group and indicates that the reduction to cerebral oxygenation is driven by this reduction in cerebral perfusion.

### Transient hyperoxia had region-specific effects on mitochondrial respiration and bioenergetics

Both the standard oxygen and air groups had significantly decreased striatal mitochondrial Complex I- (CI), combined CI&CII, and CIV-driven respiration, and reduced maximal electron transfer system capacity relative to the rapid-wean group (allp< 0.05, Figure 3A). CII-driven respiration in the standard group was significantly lower than in the rapid-wean group (*P* < 0.05). No effect of oxygen exposure on the remaining outcome measures were observed, or when data was expressed relative to ETS capacity (all p> 0.05, Figure 5A). In the cortex (Figure 4 and Figure 5B), only LEAK respiration relative to ETS capacity was reduced in the standard and air groups relative to the rapid-wean group (*p* < 0.05).

**Figure 3.**
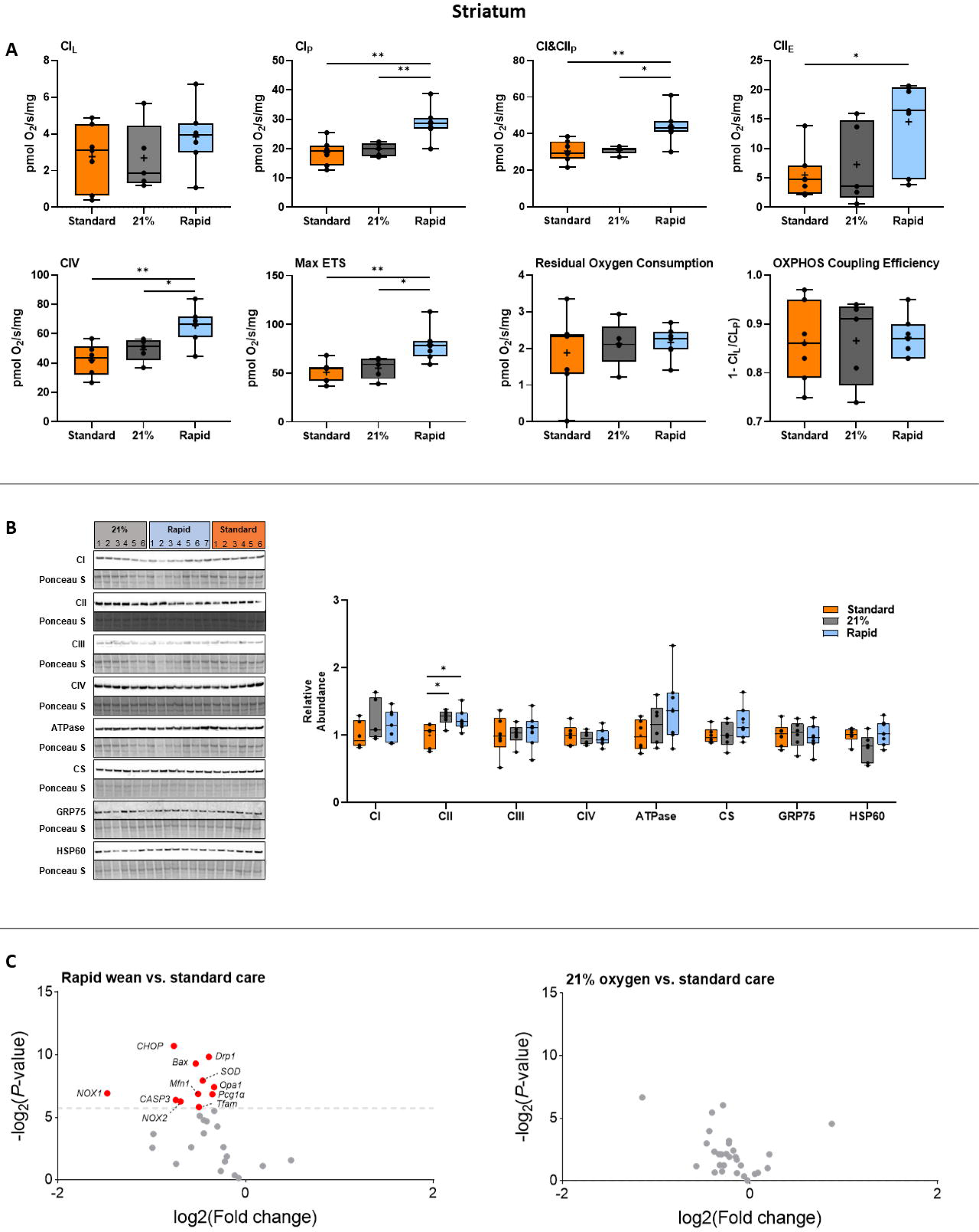
Integrated analysis of the striatum following severe perinatal asphyxia and different re-oxygenation strategies. **(A)** Mitochondrial respiration rates in the striatum for lambs in the standard oxygen treatment group (“Standard”, orange), rapid-wean group (“Rapid”, blue), and air group (“21%”, black) for leak respiration (CI_L_), Complex I-linked respiration (CI_P_), CI&CII-linked respiration (CI&CII_P_) and CII-linked respiration (CII_E_), Complex IV activity, maximal electron transfer (ETS) capacity (Max ETS), Residual Oxygen Consumption, oxidative phosphorylation (OXPHOS) coupling efficiency (OXPHOS Efficiency). **(B)** Protein abundance of electron transfer system complexes (Complex I [CI], Complex II [CII], Complex III [CIII], Complex IV [CIV], ATP synthase [ATPase]), citrate synthase (CS), 75-kDa glucose-regulated protein (GRP75), and 60-kDa heat shock protein (HSP60). Note: some proteins were probed on the same western blot (CI/CIII/ATPase and GRP75/HSP60). (**C)** Volcano plot visualizations of key genes involved in mitochondrial biogenesis, apoptosis and ER stress-induced apoptosis, antioxidant defense, and in the generation of free radicals were quantified in the rapid-wean and air groups compared to standard care. Volcano plots show fold change [log2(fold change); i.e., log2(fold change = 2) = 1; x-axis] and all gene expression changes [−log2(adjusted P-value); y axis]. The grey dashed line indicates the threshold of significance (adjusted P<0.05, using Benjamini- Hochberg correction for multiple comparisons). Genes that were downregulated are presented in red. Asterisks in black represent statistically significant differences between the rapid-wean versus standard oxygen group, and rapid-wean versus air groups, respectively (n = 5- 7/group). * p <0.05, ** p <0.01; t-test, Mann-Whitney U test, or one-way ANOVA, with Tukey post hoc tests as appropriate.

**Figure 4.**
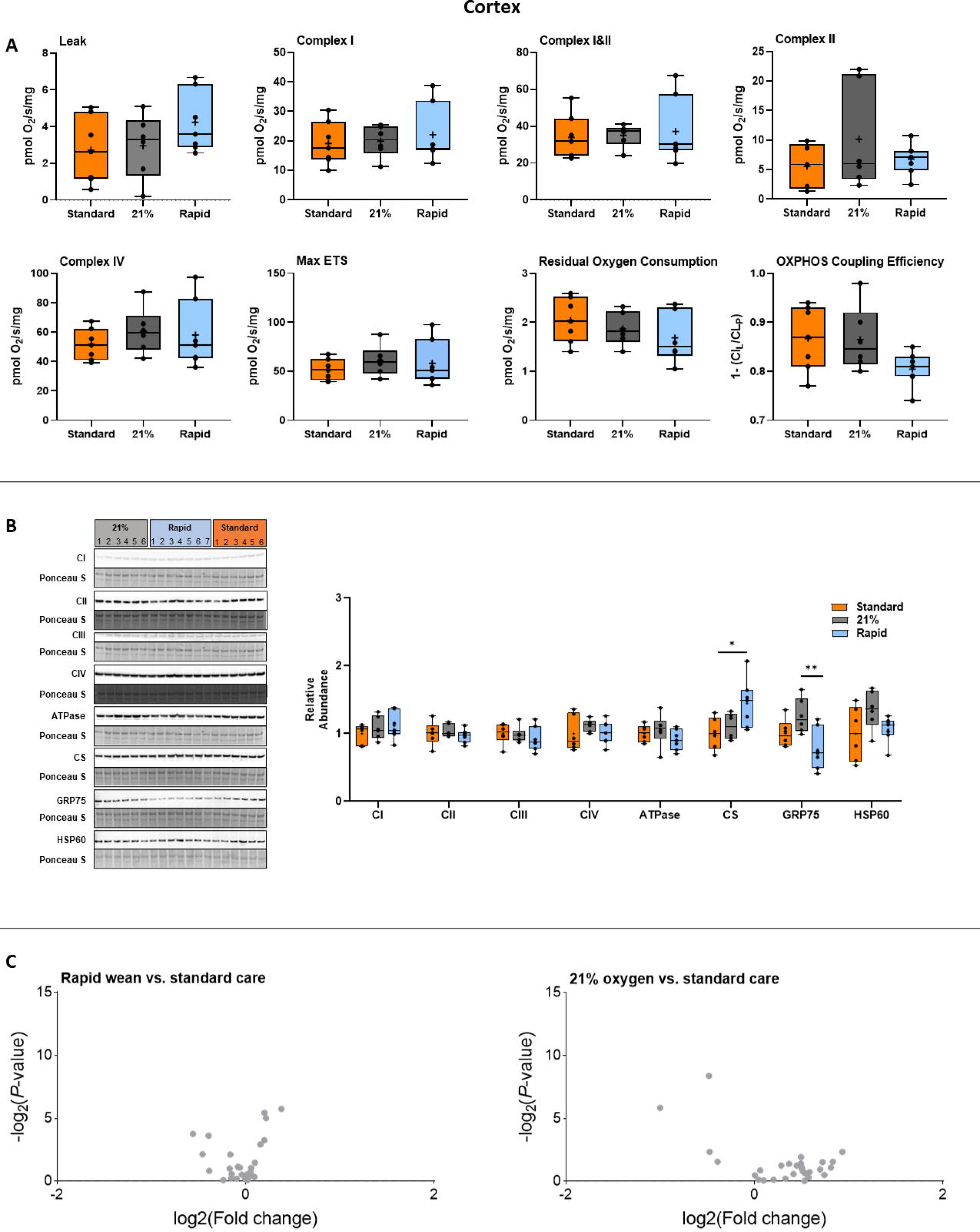
Integrated analysis of the cortex following severe perinatal asphyxia and different re-oxygenation strategies. **(A)** Mitochondrial respiration rates in the cerebral cortex for lambs in the standard oxygen treatment group (“Standard”, orange), rapid-wean group (“Rapid”, blue), and air group (“21%”, black) for leak respiration (CI_L_), Complex I-linked respiration (CI_P_), CI&CII-linked respiration (CI&CII_P_) and CII-linked respiration (CII_E_), Complex IV activity, maximal electron transfer (ETS) capacity (Max ETS), Residual Oxygen Consumption, oxidative phosphorylation (OXPHOS) coupling efficiency (OXPHOS Efficiency). (**B**) Protein abundance of electron transfer system complexes (Complex I [CI], Complex II [CII], Complex III [CIII], Complex IV [CIV], ATP synthase [ATPase]), citrate synthase (CS), 75-kDa glucose-regulated protein (GRP75), and 60-kDa heat shock protein (HSP60). Note: some proteins were probed on the same western blot (CI/CIII/ATPase and GRP75/CS). (**C)** Volcano plot visualizations of key genes involved in mitochondrial biogenesis, apoptosis and ER stress-induced apoptosis, antioxidant defence, and in the generation of free radicals were quantified in the rapid-wean and air groups compared to standard care. Volcano plots show fold change [log2(fold change); i.e., log2(fold change = 2) = 1; x-axis] and all gene expression changes [−log2(adjusted P-value); y axis]. No differences in gene expression were observed with the Benjamini-Hochberg correction for multiple comparisons. Asterisks in black represent statistically significant differences between the rapid-wean versus standard oxygen group, and rapid-wean versus air groups, respectively (*n* = 5-7/group). * p <0.05, ** p <0.01; t-test, Mann-Whitney U test, or one-way ANOVA, with Tukey post hoc tests as appropriate.

**Figure 5.**
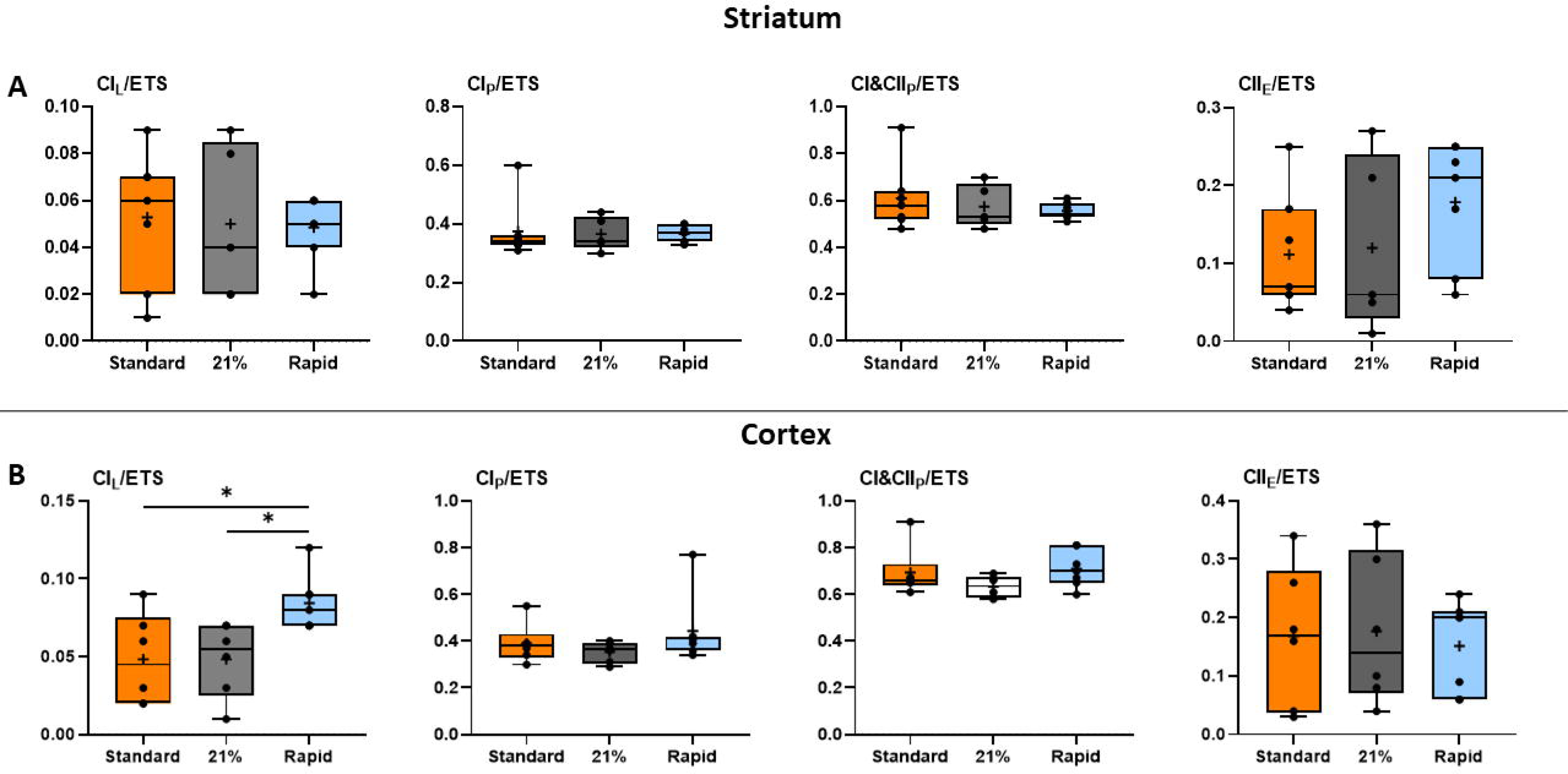
Mitochondrial respiration rates normalized to maximal electron transfer (ETS) capacity. Relative respiration rates in the striatum **(A)** and cerebral cortex **(B)** for lambs in the standard oxygen treatment group (“Standard”, orange), rapid-wean group (“Rapid”, blue), and air group (“21%”, black) for leak respiration (CIL/ETS), Complex I-linked respiration (CIP/ETS), CI&CII-linked respiration (CI&CIIP/ETS) and CII-linked respiration (CIIE/ETS).

Striatal abundance of CII was increased in both the air and rapid-wean groups relative to the standard oxygen group (*p* < 0.05, Figure 3B). The abundances of the remaining complexes were not altered in the striatum or cortex (*p* > 0.05, Figure 3B and Figure 4). In the cortex, the abundance of citrate synthase, a proxy marker for mitochondrial content, and 75-kDa glucose-regulated protein (GRP75), a mitochondrial stress protein, were increased and decreased, respectively in the rapid-wean group relative to the standard care and air groups (both p < 0.05, Figure 4).

Relative to the standard oxygen group, rapid-wean resulted in significant downregulation of multiple genes in the striatum (Figures 3C, Figure 4C), including genes that encode for NADPH oxidase enzymes which are crucial in the generation of free radicals (*NADPH Oxidase 1* and *2*, *Nox1* and *Nox2*), as well as those involved in mitochondrial biogenesis (*Tfam, Opa1, Drp1, Mfn1, PGC-1α*), apoptosis and ER stress-induced apoptosis (*Bax, Casp3*, and *CHOP*), and antioxidant defence (*SOD2* and catalase). No gene expression changes were found in the striatum in the air cohort relative to standard oxygen, nor in the cerebral cortex.

## Discussion

Our findings indicate that tissue hyperoxia occurs with brief oxygen supplementation after ROSC, is driven by rapid and prolonged increase in cerebral blood flow, and is particularly marked in the striatum. Tissue hyperoxia impairs mitochondrial respiration during the early latent phase in a region-specific manner. Using 100% oxygen during CPR but rapidly weaning oxygen to air following ROSC prevents cerebral overperfusion, hyperoxia and improves mitochondrial respiratory function.

Recent lamb studies found reductions in global cerebral oxygen delivery, as well as lower blood oxidized/reduced glutathione ratio, in lambs that had an abrupt wean in FiO_2_ from 100% to air following ROSC compared to a gradual wean.^6,7^ Our results support and extend these findings, providing evidence of marked hyperoxia at the cerebral tissue level in both the striatum and the cerebral cortex, with corresponding impairment of mitochondrial function. These findings provide a biological basis for our recently reported observation that early hyperoxemia was causally related to an increased risk of death or major disability among infants with hypoxic-ischemic encephalopathy enrolled in the Infant Cooling Evaluation trial.^22^

The effects of transient hyperoxia may be especially stark in the context of severe asphyxia, where there are currently no clinical studies prospectively evaluating oxygen supplementation strategies.^8,9^ Randomized trials of newborns with mild perinatal hypoxic-ischemia who received respiratory support without chest compressions corroborated the findings of animal models, finding lower mortality when resuscitation commenced in air versus 100% oxygen, informing current recommendations.^17,23–25^ Pre-clinical studies in non-transitioning, severely asphyxiated piglets evaluated a prolonged period (10-30 min) of exposure to 100% oxygen.^26–28^ This duration of hyperoxic exposure is now uncommon in advanced neonatal care settings. Our standard oxygen arm which provided 100% oxygen for 5 min, is more representative of the current spectrum of practice. While some practitioners may wean oxygen more quickly, we estimated that this timeframe mimics the time usually taken to first determine ROSC, obtain a reliable pulse-oximetry signal, and subsequently wean oxygen.

Two previous studies evaluated shorter periods of 100% oxygen use. Perez-de-sa *et al.*^29^ limited 100% oxygen to 3 minutes from the start of ventilation in less asphyxiated lambs that mostly did not require CPR. Linner *et al.*^30^ limited 100% oxygen to 3 minutes from commencing CPR in non-transitioning piglets. Consistent with our findings, both studies report that even this brief exposure was insufficient to prevent cerebral hyperoxia. Markers of injury or oxidative stress were not reported. Importantly, we showed that the hyperoxia was primarily driven by cerebral overperfusion, as evident by significantly elevated carotid blood flow in the standard care group compared to the rapid wean and air groups. The cerebral overperfusion is driven by a rapid increase in pulmonary blood flow, consequent to a rapid oxygen dependent fall in pulmonary vascular resistance, which increases cardiac output. Chandrasekharan and colleagues previously showed higher pulmonary blood flow in the minutes after birth in preterm lambs resuscitated with 100% oxygen compared to those ventilated with 21%.^31^ These findings highlight the critical relationship between oxygen, pulmonary and cerebral perfusion and cerebral hyperoxia immediately after birth.

Our data shows that using 100% oxygen until 5 minutes after ROSC, or using air and increasing oxygen from 5 minutes after ROSC, decreases both Complex I– and Complex II– driven respiration, Complex IV activity, and maximal ETS capacity in the striatum, suggesting that the oxidative machinery is not able to synthesize ATP at rates necessary for maintaining electrochemical gradients to support brain function in the first hour after asphyxia.^32^ When respiratory rates were normalized to maximal ETS capacity, they were no longer lower, suggesting that the apparent limitation of mitochondrial OXPHOS capacity observed in these two groups is related to an overall decrease in absolute respiratory rates and ETS capacity. These findings support previous studies in 4-week-old piglets following asphyxia and subsequent exposure to 100% oxygen.^13–15^ Persistent decreases in mitochondrial respiration may ultimately limit cerebral recovery, and play an important role in the development of secondary neurologic injury and contribute to long-term cognitive dysfunction.

In the striatum, genes involved in mitochondrial biogenesis (*PGC-1α, Tfam*), fusion (*Mfn1 and Opa1*) and fission (*Drp1*) were decreased in the rapid-wean group relative to the standard care group, while mitochondrial Complex II protein abundance was higher. These changes may serve as a mechanism to stabilize existing mitochondrial structures and dampen oxidative stress produced by damaged mitochondria following asphyxia. The downregulation of the NADPH oxidases, major sources of ROS, and key genes involved in apoptosis and ER-stress-induced cell death (*Bax, Casp3* and *CHOP*), supports this hypothesis. The increase in Complex II protein abundance points towards an increased tricarboxylic acid (TCA) capacity, and could further serve as a mechanism to help maintain mitochondrial function and sustain ATP production following the asphyxic insult.

Cortical Complex I LEAK to OXPHOS capacity was found to be increased in the rapid-wean group relative to the standard care and air groups. Additionally, there was a decrease in the abundance of GRP75, which is crucial for mitochondrial homeostasis and stress response, while citrate synthase, a proxy for mitochondrial content, was increased. These changes likely reflect compensatory mechanisms to maintain mitochondrial respiration and reduce mitochondrial stress following asphyxia. These results contrast previous studies in 4-week-old piglets, which reported significant decreases in Complex I and Complex II-driven respiration in the cortex following asphyxia and CPR.^14,15^ These region-specific differences in mitochondrial respiratory capacity and bioenergetics between studies likely reflect differences in cortical maturation, procedural variables such as duration of asphyxia, percent of inspired oxygen provided during CPR, location of tissue extraction, and time post-ROSC of tissue collection.

Our study has some limitations. We used a model of acute-severe asphyxia induced by complete umbilical cord occlusion. The etiology of severe asphyxia in human newborns is varied and may be caused by other acute, or partial, prolonged insults. The cerebrovascular responses following ROSC may be different under these circumstances. Some infants with severe asphyxia may have lung disease such as meconium aspiration syndrome. Supplemental oxygen is likely to assist in the respiratory transition in these instances, but the balance of risks with regard to cerebral hyperoxia is unknown. The major strengths include the use of a well-established large-animal model with similar transitional physiology to humans, and instrumentation to measure both regional cerebral tissue oxygenation and corresponding mitochondrial function.^9,33^

Our findings indicate that interventions to prevent hyperoxia-mediated brain injury during resuscitation of neonates following acute-profound asphyxia should focus on the period following ROSC. The risk of cerebral hyperoxia occurs not during CPR, but rather during the post-asphyxial overshoot in cerebral blood flow. A “rapid-wean” strategy facilitates the maximizing of blood oxygen content to achieve ROSC,^34^ while negating the risk of subsequent cerebral tissue hyperoxia and mitochondrial injury. With this strategy, brief exposure to supplemental oxygen in the seconds after ROSC prior to weaning may reduce the need for later supplemental oxygen (as seen in the 21% group), preventing delayed hyperoxia. Further work is needed to evaluate the effect of transient hyperoxia in partial, prolonged asphyxia, its interaction with therapeutic hypothermia, and in preterm asphyxia.^35,36^ Strategies to better target cerebral oxygenation and limit hyperoxia-mediated brain injury are needed.

## Acknowledgements

We thank the staff at the Monash Animal Research Platform for their contribution to this research.

## Availability of data and materials

The datasets used and/or analyzed during the current study are available from the corresponding author on reasonable request.

## Supplementary material

Supplementary material with further details on study methods are available online.

