## Supplementary material for "Rapid oxygen titration following cardiopulmonary resuscitation mitigates cerebral overperfusion and striatal mitochondrial dysfunction in asphyxiated newborn lambs"

**Supplementary Materials**

**Supplementary Methods**

Pregnant Border-Leicester ewes (Ovis aries) were housed in floor pens with access to food and water ad libitum and were monitored daily for health and well-being 7 days prior to experiments. All experiments were carried out between 8am and 5pm in a dedicated non-sterile animal laboratory operating facility. At 139±2 days’ gestation (mean ± SD; term ~148 days), ewes were anaesthetized by intravenous injection of thiopentone sodium (20 mg/ kg; Jurox, NSW, Australia), followed by tracheal intubation and delivery of inhaled anaesthesia (isoflurane 1.5%–2.5% in oxygenated air; Bomac Animal Health, NSW, Australia). The oxygen concentration of the carrier gas was titrated between 21-80% FiO_2_ to maintain the ewes SpO_2_ between 90-95%.

**Instrumentation**

An ultrasonic flow transducer of appropriate size (Transonic Systems, Ithaca, NY, USA) was placed around the left carotid artery and left main pulmonary artery, accessed via a left thoracotomy. A pressure transducer (PD10; DTX Plus Transducer; Becton Dickinson, Singapore) was used to measure blood pressure directly within the left brachial artery. Catheters were inserted into the right jugular vein (occlusive) and right carotid artery (non-occlusive) for central venous and pre-ductal arterial blood gas sampling respectively. After closure of the incisions in the neck and chest, the fetal trachea was intubated with a 4.5 mm cuffed endotracheal tube and lung liquid was passively drained. A transcutaneous arterial oxygen saturation probe (Masimo SET Pulse Oximeter Sensor LNCS Neo-3, Irvine CA, USA) was placed around the right forelimb of the lamb.

A fibre-optic oxygen probe (oxygen bare-fibre sensor NX-BF/O/E, Oxford Optronics, Abingdon, United Kingdom) was placed in the right carotid artery and the right cerebral striatum. For the latter, a single burr hole (2 mm diameter) was made 15 mm anterior and 5 mm lateral to bregma (the point at which the coronal and sagittal sutures of the skull meet) and a 20G intracath was introduced to a depth of 20mm. The tip of the oxygen probe was placed just outside the end of the intracath to measure tissue pO_2_ in the caudate nucleus. A Near Infrared Spectroscopy optode (Casmed Foresight, CAS Medical Systems, Branford, CT, USA) was placed on the shaved scalp over the left frontal cortex, secured using opaque material and used to continuously measure cerebral tissue oxygen saturation. A rectal probe was inserted to monitor temperature.

Following instrumentation, the fetus was completely exteriorized, physiological parameters were allowed to stabilize and asphyxia subsequently induced by cord clamping while withholding ventilation. Lambs were randomly assigned using a web-based random sequence generator (www.random.org/lists) to one of the 3 oxygen treatment groups to minimize confounding. Group allocation was unblinded to the investigators.

**Management after birth**

Following ROSC, lambs received ventilator-driven breaths with warm, humidified gases using peak inspiratory pressure of 30 cmH_2_O, end-expiratory pressure of 5 cmH_2_O and rate 60 ventilations per minute (Babylog 8000+, Dräger, Lübeck, Germany). From 10 minutes after ROSC lambs were switched to volume-guarantee mode (7 mL/kg). From 5 minutes after ROSC, FiO_2_ was adjusted in all 3 groups to target arterial oxygen saturation between 90% and 95%. Ventilation was adjusted to maintain PaCO_2_ between 45 and 55 mmHg. Ventilator changes to achieve oxygen saturation and PaCO_2_ targets were performed by the same 2 neonatologists (SB and CTR) for all lambs unblinded to the group allocation. Lambs were sedated from 10 minutes after ROSC (Alfaxalone intravenous 5–15 mg/kg/hour in 5% dextrose; Jurox) to prevent spontaneous breathing.

**Data acquisition and analysis**

Continuous digital recordings of physiological parameters were made in real-time (1kHz) using a data acquisition system (PowerLab; ADInstruments, NSW, Australia). Paired carotid arterial and jugular venous blood gas parameters (temperature-adjusted) were measured using a blood gas analyzer (ABL90 Flex Plus, Radiometer, Copenhagen, Denmark). The first was taken before induction of asphyxia, at terminal asphyxia, during CPR and prior to administration of adrenaline (n= 6, arterial only), at ROSC, and then at 3, 6, 9, 12, 15, 20, 25, 30, 40, 50 and 60 minutes after ROSC.

**Tissue collection and analysis**

At 1 hour after ROSC, lambs were then euthanized (sodium pentobarbitone >100 mg/kg, intravenous) and major organs were excised and weighed. Fetal brains were hemisected; fresh tissue samples (~10 mg) from the striatum (at the level of where the oxygen probe was placed) and cerebral cortex (at the level of the ansate sulcus) were collected from the left hemisphere and placed in ice-cold buffer (MiR06). The remaining portions of the left hemisphere were frozen in liquid nitrogen and stored at −80 ◦C for subsequent molecular analysis. The right hemisphere was immersion-fixed in 4% paraformaldehyde (PFA) for histopathological analysis.

**High-resolution respirometry**

Cerebral mitochondrial respiratory rates were measured by high-resolution respirometry as previously described.^1,2^ Fresh biopsies (~10 mg) obtained from the striatum and cerebral cortex (*n* = 6-7/group) were homogenized in respiratory medium (MiR06) and then transferred into pre-calibrated oxygraph chambers (Oxygraph 2k, Oroboros Instruments, Austria). LEAK respiration (non-phosphorylating state of intrinsic uncoupling) was measured in the presence of malate (2 mM) and pyruvate (5 mM). ADP (10 mM; saturating concentration) was added to stimulate OXPHOS, followed by glutamate (10 mM) to drive OXPHOS capacity via Complex I (CIP). Maximal electron flux through Complex I and Complex II was achieved through the addition of succinate (10 mM, CI&CIIP). Subsequently, inhibition of CI by rotenone (0.5 µM) provided measurement of CII-linked OXPHOS capacity (CIIP). Maximal non-coupled electron transfer system (ETS)-respiratory capacity was measured with trifluoromethoxy carbonyl-cyanide phenylhydrazone (FCCP, titration doses at 0.5 µM). Antimycin A (2.5 µM), an inhibitor of Complex III, provided a measure of residual (non-mitochondrial and mitochondrial) oxygen consumption. Subsequently, inhibition of CI by rotenone (0.5 µM) provided measurement of CII-linked OXPHOS capacity (CIIP). Cytochrome c (10 µM) was added to check the integrity of the mitochondrial membranes with data excluded if respiration increased by >15%. Respiratory capacities were also normalized to maximal non-coupled ETS capacity. OXPHOS coupling efficiency was calculated to represent the net OXPHOS capacity, corrected for leak respiration (1- Leak/CIP).

**Western blot analyses**

The abundance of mitochondrial regulatory proteins of the ETS, and mitochondrial- and oxidative-stress-related proteins, were determined in frozen brain samples using western blotting (*n* = 6-7 group) as previously described.^1,2^ Total protein was extracted from tissue samples (~30 mg) using RIPA buffer (Thermo Fisher Scientific, Waltham MA, U.S.A), and lysate protein concentrations determined using a BCA assay (Sigma-Aldrich, St. Louis, U.S.A.) according to the manufacturer's protocol. Equivalent amounts of protein (20 μg) were separated by electrophoresis on a 4-12% polyacrylamide gel (Thermo Fisher Scientific, Waltham MA, U.S.A). Protein was transferred to a nitrocellulose membrane, then stained with Ponceau-S to normalize for protein loading. Membranes were probed overnight at 4°C with the following primary antibodies: mitochondrial stress marker glucose-regulated protein 75 (GRP75, Thermo Fisher Scientific, Waltham MA, U.S.A; 1:3000; RRID: AB_2120465), heat shock protein 60 (HSP60)- localized in the mitochondria and essential in the integrity and function of the mitochondrial respiratory transfer system (Thermo Fisher Scientific, Waltham MA, U.S.A; 1:2000; catalogue number: 3329-RBM6-P1ABX), citrate synthase (CS)-a proxy marker for mitochondrial content (Abcam, Cambridge, UK; 1:1000; RRID: RRID:AB_10678258), ETS Complexe II (Thermo Fisher Scientific, Waltham MA, U.S.A; 1:1000; RRID: AB_2532233) and Complex IV (Thermo Fisher Scientific, Waltham MA, U.S.A; 1:1000; RRID: AB_2532240), and an antibody cocktail for the remaining ETS complexes (OXPHOS antibody cocktail; Thermo Fisher Scientific, Waltham MA, U.S.A; 1:1000; RRID: AB_2533835) overnight followed by HRP-linked secondary antibodies (Thermo Fisher Scientific, Waltham MA, U.S.A and Aglient, Santa Clara, CA, U.S.A.; both 1:5000; RRID:AB_325993 and RRID: AB_2728719). Protein bands were visualized using enhanced chemiluminescence (Thermo Fisher Scientific, Waltham MA, U.S.A) and the iBright imaging system (Thermo Fisher Scientific, Waltham MA, U.S.A) and then quantified using ImageJ (NIH).

**Gene expression analyses**

RNA was extracted from frozen brain tissue (~30 mg, RNeasy Plus Mini Kit, Qiagen; Thermo Fisher Scientific, Waltham MA, U.S.A) and yields assessed using a Nanodrop (Nanodrop, Analytical Technologies, Biolab, Concord, Ontario, Canada). Reverse transcription of the extracted mRNA was performed using the High-Capacity cDNA Reverse Transcription Kit (Thermo Fisher Scientific, Waltham MA, U.S.A). Gene expression was analyzed using a Fluidigm Dynamic array Biomark HD system (Fluidigm, San Francisco, CA, U.S.A). Genes involved in mitochondrial biogenesis, oxidative stress, apoptosis and ER stress-induced apoptosis (Bax, Casp3, and CHOP), and antioxidant defence (SOD2 and catalase) were analyzed in both the striatum and cerebral cortex (Supplementary Table S1). Levels of mRNA expression relative to the housekeeping gene which was the most stable between the different oxygen support groups (YWHAZ) were determined using the 2–ΔΔCT method. The results were expressed as fold changes relative to the standard care group.

**Supplementary Table**

| **Gene ID** | **Taqman Assay ID** |
| --- | --- |
| NRF1 | Oa04660399_m1 |
| NRF2 | Oa04815261_g1 |
| TFAM | Oa03260079_g1 |
| PGC-1α | Oa01208835_m1 |
| PPARγ | Oa04658385_m1 |
| UCP2 | Oa03225185_g1 |
| ANT1 | Oa04655658_g1 |
| Drp1 | Oa04916359_m1 |
| Fis1 | Oa03227775_m1 |
| Opa1 | Oa04313744_m1 |
| MFN2 | Oa04843335_g1 |
| HSP10 | Bt03213904_m1 |
| HSP60 | Oa04923161_g1 |
| ClpP | Oa03239784_g1 |
| mtDNA | Bt03218015_g1 |
| CHOP | Oa04654052_g1 |
| MFN1 | Oa04295229_m1 |
| c-MYC | Oa04658354_m1 |
| HIF-1α | Oa04877335_m1 |
| MPO | Oa04654413_g1 |
| NOX1 | Oa04709255_g1 |
| NOX2 | Oa04793417_m1 |
| GPX1 | Oa04911462_g1 |
| CAT | Oa03228713_m1 |
| SOD2 | Oa04657474_m1 |
| CASP3 | Oa04817361_m1 |
| BAX | Oa03211776_g1 |
| BAK1 | Oa04906761_m1 |
| BCL2 | Oa04888158_m1 |
| 18S (Housekeeping) | Oa4906333_g1 |
| B2M (Housekeeping) | Oa04900279_Mh |
| YWHAZ (Housekeeping) | Oa04913608_m1 |

**Supplementary Table S1. Genes of interest**
